# *C. elegans* HIF-1 is broadly required for survival in hydrogen sulfide

**DOI:** 10.1101/174581

**Authors:** Irini Topalidou, Dana L Miller

**Affiliations:** University of Washington School of Medicine, Department of Biochemistry, Seattle, WA 98195

**Keywords:** hydrogen sulfide, hypoxia, tissue-specific expression

## Abstract

Hydrogen sulfide is common in the environment, and is also endogenously produced by animal cells. Although hydrogen sulfide is often toxic, exposure to low levels of hydrogen sulfide improves outcome in a variety of mammalian models of ischemia-reperfusion injury. In *Caenorhabditis elegans*, the initial transcriptional response to hydrogen sulfide depends on the *hif-1* transcription factor, and *hif-1* mutant animals die when exposed to hydrogen sulfide. In this study, we use rescue experiments to identify tissues in which *hif-1* is required to survive exposure to hydrogen sulfide. We find that expression of *hif-1* from the *unc-14* promoter is sufficient to survive hydrogen sulfide. Although *unc-14* is generally considered to be a pan-neuronal promoter, we show that it is active in many non-neuronal cells as well. Using other promoters, we show that pan-neuronal expression of *hif-1* is not sufficient to survive exposure to hydrogen sulfide. Our data suggest that *hif-1* is required in many different tissues to direct the essential response to hydrogen sulfide.

## INTRODUCTION

Hydrogen sulfide (H_2_S) in the environment is produced by industrial sources and natural sources, including volcanic deposits and anaerobic bacteria (Beauchamp *et al.*, 1984). H_2_S is also endogenously produced as a product of the cysteine biosynthesis through the transsulfuration pathway, and endogenous H_2_S has important roles in cellular signaling (Li *et al.*, 2011; Vandiver and Snyder, 2012; Wang, 2012). Chronic exposure to relatively low concentrations of environmental H2S in humans has been associated with neurological, respiratory, and cardiovascular dysfunction (Kilburn and Warshaw, 1995; Richardson, 1995; Bates *et al.*, 2002). However, transient exposure to low H_2_S has also been shown to improve outcome in many mammalian models of ischemia-reperfusion injury (Bos *et al.*, 2015; Wu *et al.*, 2015). It is possible that the biological effects of exogenous H_2_S exposure, both beneficial and detrimental, result from activation of pathways that are normally regulated by endogenous H_2_S.

*C. elegans* is an excellent system to define physiological responses to exogenous H2S. In addition to powerful genetics, all cells are directly exposed to the gaseous environment (Shen and Powell-Coffman, 2003). This feature allows for control of cellular H_2_S exposure without confounding factors from physiological regulation gas delivery. *C. elegans* grown in H_2_S are long-lived, thermotolerant, and resistant to hypoxia-induced disruptions of proteostasis (Miller and Roth, 2007; Fawcett *et al.*, 2015). HIF-1 directs the transcriptional response to H_2_S in *C. elegans* (Budde and Roth, 2010; Miller *et al.*, 2011). HIF-1 is a highly-conserved transcription factor best known for regulating the transcriptional response to low oxygen (hypoxia) in metazoans (Semenza, 2000; Semenza, 2001). *C. elegans hif-1* mutant animals are viable and fertile in room air but die if exposed to hypoxia during embryogenesis (Jiang *et al.*, 2001; Miller and Roth, 2009). In contrast, exposure to H_2_S is lethal for *hif-1* mutant animals at all developmental stages (Budde and Roth, 2010).

Several studies have argued for neuronal-specific functions of HIF-1, though the *hif-1* promoter is active in most, if not all, cells and HIF-1 protein is stabilized ubiquitously in *C. elegans* exposed to either hypoxia or H_2_S (Jiang *et al.*, 2001; Budde and Roth, 2010). Neuronal expression of *hif-1* in hypoxia is reported to be sufficient to prevent hypoxia-induced diapause and to increase lifespan through induction of intestinal expression of *fmo-2* (Miller and Roth, 2009; Leiser *et al.*, 2015). Furthermore, neuronal CYSL-1 protein regulates the activity of HIF-1 to modulate behavioral responses to changes in oxygen availability (Ma *et al.*, 2012). These data motivated us to determine if neuronal HIF-1 activity is sufficient for *C. elegans* to survive exposure to H_2_S.

In this study, we used tissue-specific rescue of *hif-1* to define the site of essential HIF-1 activity in low H_2_S. We found that expression of *hif-1* from the *unc-14* promoter was sufficient for survival in H_2_S. Although considered a pan-neuronal promoter (Ogura *et al.*, 1997; Pocock and Hobert, 2008), our data indicate that the *unc-14* promoter is also broadly expressed in non-neuronal cells. We show that *hif-1* expressed from the pan-neuronal *rab-3* promoter is not sufficient for viability in H_2_S. We further demonstrate that expression of *hif-1* in muscle, hypodermis, or intestine is not sufficient for viability in low H_2_S. Together, our data indicate that the activity of HIF-1 may be required in multiple tissues to coordinate the organismal response to H_2_S.

## MATERIALS AND METHODS

### Strains

Strains were grown at room temperature on nematode growth media plates (NGM) seeded with the OP50 strain of *E. coli* (Brenner, 1974). All strains were derived from N2 (Bristol). Full genotypes of strains used in this study are in Table 1. To sequence the *Punc-14::hif-1* junction of *otIs197*, the region was amplified with forward primer oET479 (5’-GTTGTCCACCATCACAGTAATACG) and reverse primer oET480 (5’-ACGACGGCGTTCCATG). The oET479 primer was used for sequencing.

**Table 1:**
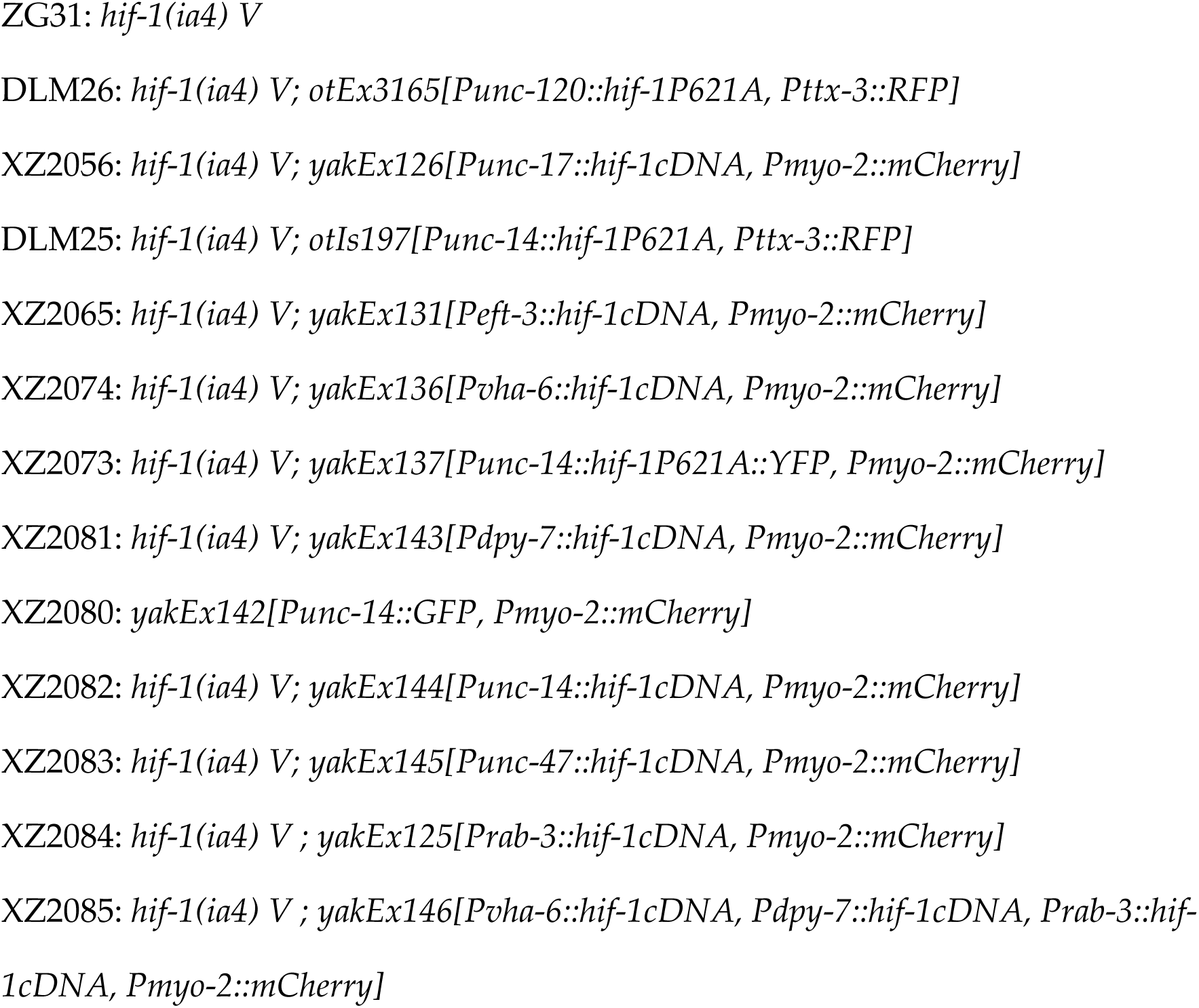
Strains used in this study.

### Constructs and transgenes

All constructs were made using the multisite Gateway system (Invitrogen) where a promoter region, a gene region (*hif-1* cDNA or GFP), and a C-terminal 3’UTR were cloned into the destination vector pCFJ150 (Frøkjaer-Jensen *et al.*, 2008). The *hif-1* A isoform was amplified from cDNA using forward primer oET467 (5’-GGGGACAAGTTTGTACAAAAAAGCAGGCTCAATGGAAGACAATCGGAAAAGAAAC) and reverse primer oET469 (5’-GGGGACCACTTTGTACAAGAAAGCTGGGTGTCAAGAGAGCATTGGAAATGGG). For the tissue-specific rescuing experiments, an operon GFP::H2B was included in the expression constructs downstream of the 3’UTR (Frøkjær-Jensen *et al.*, 2012). This resulted in expression of untagged HIF-1 protein and histone H2B fused to GFP, which allowed for confirmation of promoter expression by monitoring GFP expression. The *unc-14* promoter (1425 bp upstream of the start codon) was amplified from genomic DNA using forward primer oET520 (5’-GGGGACAACTTTGTATAGAAAAGTTGGAGAGCAGCAGCATCTCGAG) and reverse primer oET507 (5’-GGGGACTGCTTTTTTGTACAAACTTGTTTTGGTGGAAGAATTGAGGG). All plasmids constructed were verified by sequencing. Constructs used in this study are in Table 2. Extrachromosomal arrays were made by standard injection methods (Mello *et al.*, 1991) with 10-15 ng/μl of the expression vector. At least two independent lines were isolated for each construct.

**Table 2:**
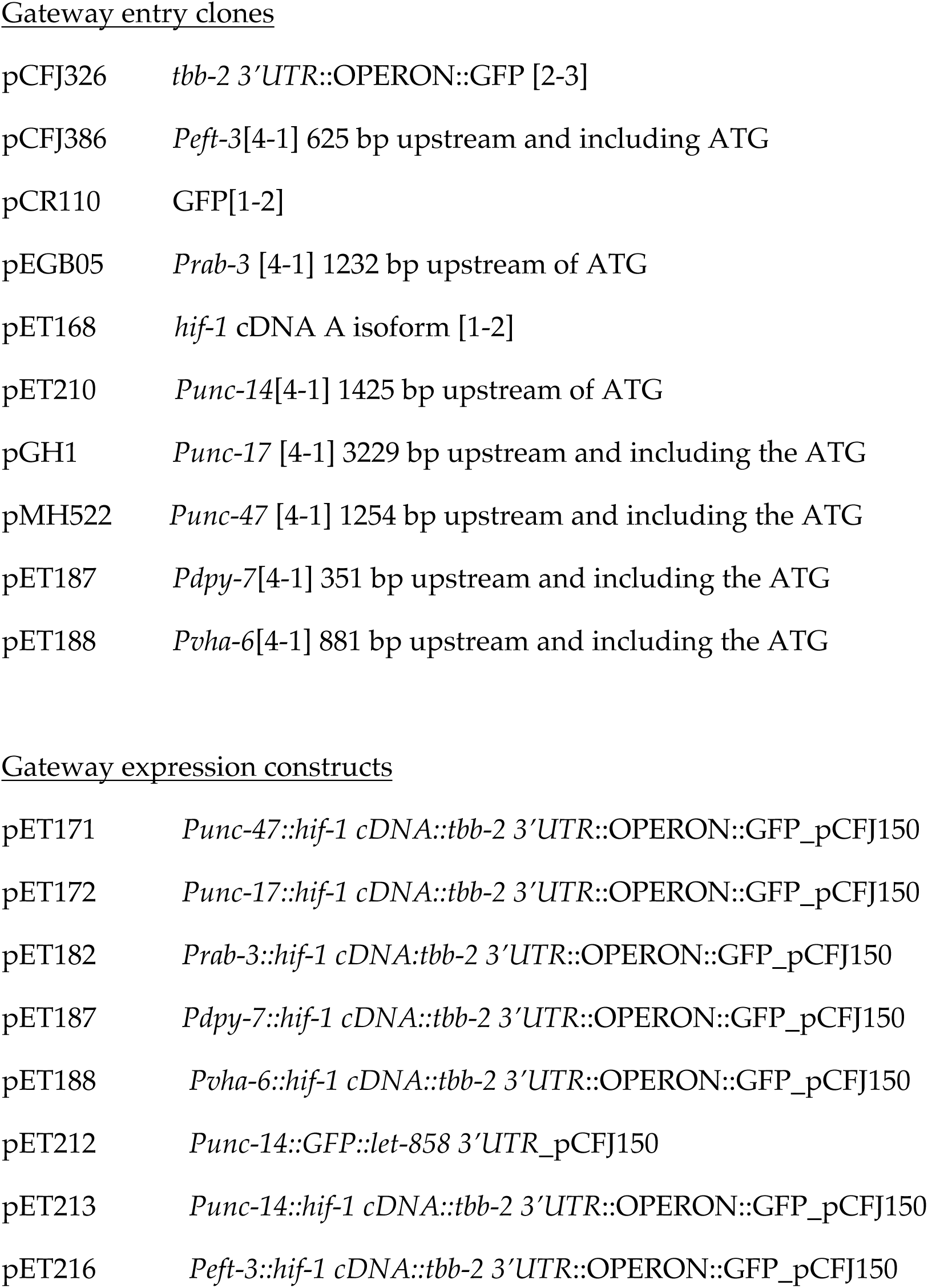
Plasmids and constructs used in this study.

Gateway entry clones
|  |  |
| --- | --- |
| pCFJ326 | *tbb-2* 3'UTR::OPERON::GFP [2-3] |
| pCFJ386 | *Peft-3*[4-1] 625 bp upstream and including ATG |
| pCR110 | GFP[1-2] |
| pEGB05 | *Prab-3* [4-1] 1232 bp upstream of ATG |
| pET168 | *hif-1* cDNA A isoform [1-2] |
| pET210 | *Punc-14*[4-1] 1425 bp upstream of ATG |
| pGH1 | *Punc-17* [4-1] 3229 bp upstream and including the ATG |
| pMH522 | *Punc-47* [4-1] 1254 bp upstream and including the ATG |
| pET187 | *Pdpy-7*[4-1] 351 bp upstream and including the ATG |
| pET188 | *Pvha-6*[4-1] 881 bp upstream and including the ATG |
| Gateway expression constructs |  |
| pET171 | *Punc-47::hif-1 cDNA::tbb-2* 3'UTR::OPERON::GFP_pCFJ150 |
| pET172 | *Punc-17::hif-1 cDNA::tbb-2* 3'UTR::OPERON::GFP_pCFJ150 |
| pET182 | *Prab-3::hif-1 cDNA::tbb-2* 3'UTR::OPERON::GFP_pCFJ150 |
| pET187 | *Pdpy-7::hif-1 cDNA::tbb-2* 3'UTR::OPERON::GFP_pCFJ150 |
| pET188 | *Pvha-6::hif-1 cDNA::tbb-2* 3'UTR::OPERON::GFP_pCFJ150 |
| pET212 | *Punc-14::GFP::let-858* 3'UTR_pCFJ150 |
| pET213 | *Punc-14::hif-1 cDNA::tbb-2* 3'UTR::OPERON::GFP_pCFJ150 |
| pET216 | *Peft-3::hif-1 cDNA::tbb-2* 3'UTR::OPERON::GFP_pCFJ150 |

### H_2_S atmospheres

Construction of atmospheric chambers was as previously described (Miller and Roth, 2007; Fawcett *et al.*, 2012). In short, H_2_S (5000 ppm with balance N_2_) was diluted continuously with room air to a final concentration of 50 ppm. Final H_2_S concentration was monitored using a custom-built H_2_S detector containing a three-electrode electrochemical Surecell H_2_S detector (Sixth Sense) as described (Miller and Roth, 2007), calibrated with 100 ppm H_2_S with balance N_2_. Compressed gas mixtures were obtained from Airgas (Radnor, PA) and certified standard to within 2% of the indicated concentration.

### Survival assays

Twenty to forty L4 animals were picked to plates seeded with OP50. Plates were exposed to 50ppm H_2_S for 20-24 hours, and then returned to room air to score viability. Death was defined as failure to move when probed with a platinum wire on the head or tail.

### Imaging

For imaging expression of GFP, larval stage 1 (L1) or first-day adult animals were mounted on 2% agarose pads and anesthetized with 50 mM sodium azide for ten minutes before placing the cover slip. The images were obtained using a Nikon 80i wide-field compound microscope.

### Reagent Availability

Strains are available upon request and have been deposited at the *Caenorhabditis* Genetics Center (cgc.umn.edu). Plasmid constructs are available upon request.

## RESULTS AND DISCUSSION

*C. elegans* requires *hif-1* to survive exposure to low H_2_S (Budde and Roth, 2010). To determine whether neuronal expression of *hif-1* was sufficient for survival in H_2_S, we used transgenic *hif-1(ia04)* mutant animals that expressed *hif-1* from heterologous promoters. We first used the available *otIs197* transgene, which expresses *hif-1* from the putative pan-neuronal *unc-14* promoter (Pocock and Hobert, 2008). We found that *hif-1(ia04); otIs197* animals survived exposure to 50 ppm H_2_S (Figure 1A). This result suggests that neuronal expression of *hif-1*, from the *unc-14* promoter, is sufficient to survive exposure to H_2_S.

**Figure 1.**
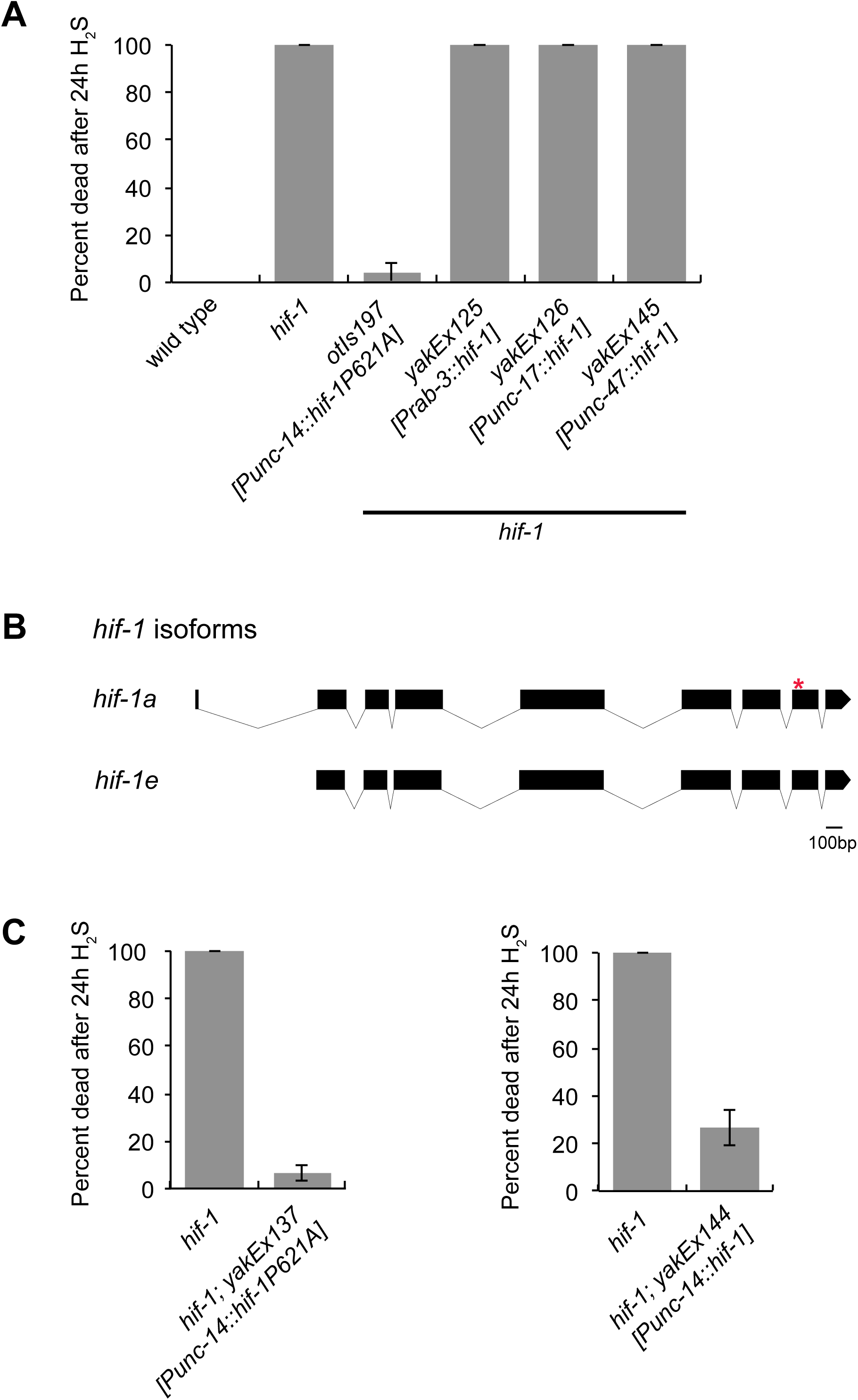
HIF-1 expression from the *unc-14*promoter rescues the H_2_S lethality of *hif-1(ia04)*mutant animals. (A) Survival of animals exposed to H_2_S. All animals have the null *hif-1(ia04)* mutation. The *otIs197* integrated array expresses a non-degradable HIF-1 variant. Other constructs were extrachromosomal arrays that express wild-type HIF-1. The *unc-14* promoter is expressed pan-neuronally (Ogura *et al.*, 1997), *rab-3* promoter is expressed in most, if not all, neurons (Nonet *et al.*, 1997), *unc-17* is expressed in cholinergic neurons (Rand *et al.*, 2000), and *unc-47* is expressed in GABAergic neurons (Eastman *et al.*, 1999). Animals were exposed to 50 ppm H_2_S starting at L4. (B) HIF-1 gene structure and predicted A and E isoforms (Wormbase, 2017). The P621A mutation that prevents degradation of *hif-1* included in *otIs197* is marked with *. (C) Survival of animals expressing HIF-1 from *unc-14* promoter exposed to H_2_S. All animals have the null *hif-1(ia04)* mutation. Expression of HIF-1 was from extrachromosomal arrays. The *yakEx137* array expresses non-degradable HIF-1(P621A) and the *yakEx144* array expresses wild-type *hif-1*. For all panels animals were exposed to 50 ppm H_2_S starting at L4. Average of three independent experiments is shown, each with n = 20-40 animals. Error bars are standard error of the mean (SEM).

To further dissect which neuronal cell type(s) HIF-1 activity was required to survive exposure to H_2_S, we generated transgenic animals that expressed *hif-1* cDNA under the control of promoters active in specific neuronal subtypes. We found that expression in neither cholinergic neurons (*Punc-17*) nor in GABAergic neurons (*Punc-47*) was sufficient to rescue the lethality of *hif-1(ia04)* mutant animals exposed to H_2_S (Figure 1A). Curiously, we also observed that expression of *hif-1* cDNA from the pan-neuronal *rab-3* promoter did not rescue survival of the *hif-1(ia04)* mutant animals (Figure 1A). This was unexpected, as expression of HIF-1 from the *unc-14* promoter (the *otIs197* transgene) was sufficient for survival in H_2_S. We therefore pursued the source of this discrepancy.

We first sought to verify the molecular nature of the *otIs197* integrated transgene. We used PCR to amplify a region from the *unc-14* promoter and the *hif-1* coding region from the *otIs197* transgenic animals. As expected, this reaction generated a single band of ∼500 bp. However, when we sequenced the resulting PCR product we discovered an insertion of an extra G immediately following the ATG of the *hif-1* cDNA. This insertion causes a frame-shift and results in a stop codon after 13 amino acids. However, the *otIs197* transgene must express some HIF-1 protein, as it can rescue many phenotypes of *hif-1* mutant animals (Pocock and Hobert, 2008; Miller and Roth, 2009; Ma *et al.*, 2012; Leiser *et al.*, 2015). The *otIs197* transgene was constructed to express isoform A of *hif-1*, though there are six predicted isoforms (Wormbase, 2017). We noted that the ATG for isoform E is approximately 20 bp downstream of the original ATG in the *hif-1* cDNA. Thus, it could be that expression of the *hif-1e* isoform is the basis of the activity of the *otIs197* transgene. Because our *Prab-3::hif-1* transgene expressed the *hif-1a* isoform, it was possible that the differences we observed from *otIs197* was due to the expression of different *hif-1* isoforms. To test this possibility we created transgenic strains expressing *hif-1a* under control of the *unc-14* promoter using a *Punc-14*::*hif-1a(P621A)::YFP* plasmid (Pocock and Hobert, 2008), which we verified had had the expected *hif-1a(P621A)* sequence. We injected this plasmid into *hif-1(ia04)* mutant animals to generate the *yakEx137* transgene. If the rescue we observed in *otIs197* was due to expression of *hif1e* rather than *hif1a*, then the animals expressing *Punc-14*::*hif-1a(P621A)::YFP* would die in H_2_S. However, these animals survived exposure to H_2_S (Figure 1C), indicating that potential expression of different isoforms did not underlie differences in survival of exposure to H_2_S.

The HIF-1 protein expressed by the *otIs197* transgene has a P621A mutation that prevents it from being hydroxylated and degraded by the proteasome (Pocock and Hobert, 2008). In contrast, the constructs we generated produced wild-type HIF-1 protein. We did not expect this feature to be salient for our experiments, since HIF-1 protein is stabilized in H_2_S due to inhibition of the hydroxylation reaction (Budde and Roth, 2010; Ma *et al.*, 2012). However, it is possible that constitutive stabilization of HIF-1 protein in neurons promotes survival in H_2_S. To evaluate this possibility, we cloned wild type *hif-1* cDNA under control of the *unc-14* promoter, including 1.4 kb upstream of the transcription start site (Ogura *et al.*, 1997). We found that *hif-1(ia04); Punc-14::hif-1* (*yakEx144*) animals survived exposure to H_2_S, similar to *hif-1(ia04); otIs197* animals (Figure 1C). We conclude that the P621A mutation in *otIs197* does not underlie the difference in survival in H_2_S that we observe for animals expressing *hif-1* from *rab-3* and *unc-14* promoters.

Given that the only other notable difference between the *Prab-3::hif-1* and *Punc-14::hif-1* constructs is the promoter elements, we hypothesized that differences between either the levels of expression from these promoters or the identity of the cells where these promoters are expressed should account for their different behavior. The transgenic constructs we generated all included an operon GFP::H2B downstream of the 3’UTR (Frøkjær-Jensen *et al.*, 2012). This resulted in expression of untagged HIF-1 protein as well as GFP::H2B. We therefore visualized GFP expression to evaluate the expression levels and cellular patterns of promoter activity. As expected, GFP expression from adult *hif-1(ia04); Prab-3::hif-1::operon::GFP::H2B* was exclusively in neurons (Figure 2A). However, when we imaged adult *hif-1(ia04); Punc-14::hif-1::operon::GFP::H2B* animals that had survived exposure to H_2_S we observed GFP expression in neurons, as expected, but also in intestinal and hypodermal cells (Figure 2B). We saw similar expression in animals that had not been exposed to H_2_S. To corroborate this observation, we cloned the *unc-14* promoter upstream of GFP and injected it into wild-type animals. We then imaged larvae (Figure 2C) and adult animals (Figure 2D) from three separate lines. We observed expression of GFP was expressed in numerous cells other than neurons including intestine, hypodermis, muscle, and the uterus. Every animal that we imaged had expression in at least one other cell type other than neurons (n = 50).

**Figure 2.**
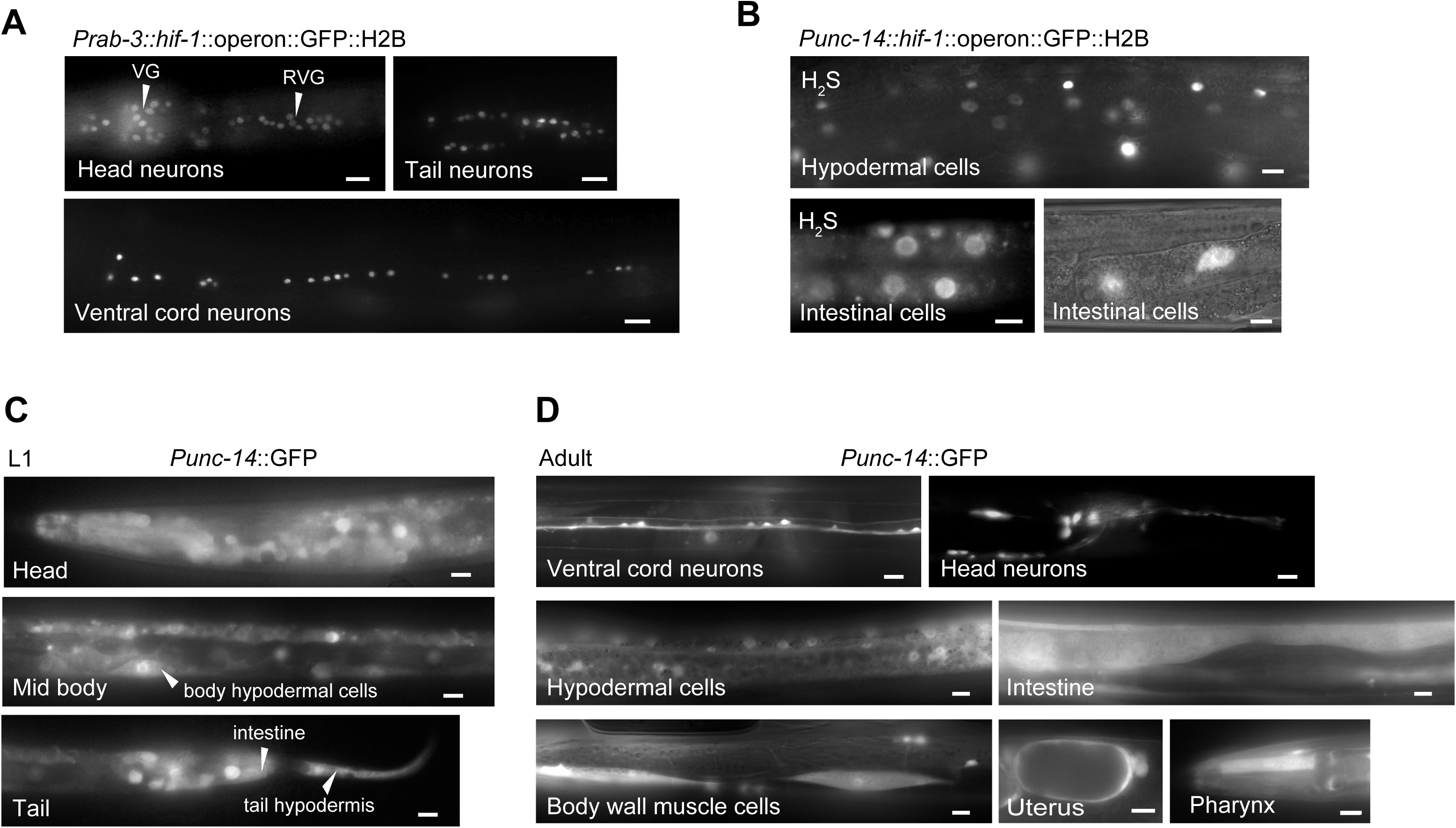
The *unc-14* promoter is active in many non-neuronal cells. (A) Visualization of GFP expressed from *Prab-3::hif-1::operon::GFP::H2B* (transgene *yak125*). Tail, head, and ventral cord neurons are shown of the ventral aspect of the same animal. VG = ventral ganglia. RVG = retrovesicular ganglia. In all images scale bar is 10 μl. (B) Representative images of adult *hif-1(ia04); Punc-14::hif-1::operon::GFP::H2B* (transgene *yakEx144*) animals. GFP expression in hypodermal and intestinal cells is shown. Scale bar is 10 μl. (C,D) Representative images of (C) L1 and (D) adult transgenic animals expressing *Punc-14::GFP* (transgene *yakEx142*). Representative animals are shown with GFP expression in hypodermis, intestine, muscle, uterus, pharynx, and neurons. Scale bars are 5μm in (C) and 10μl in (D).

Based on our understanding of *Punc-14* expression and the fact that *hif-1(ia04); Prab-3::hif-1* animals die when exposed to H_2_S (Figure 1A), we inferred that neuronal HIF-1 activity is not sufficient for survival in H2S. We therefore explored whether expression of *hif-1* exclusively in non-neuronal tissues was sufficient for survival in H_2_S. For these experiments, we generated transgenes with *hif-1* expressed under control of the *unc-120* promoter, which is active in body-wall and vulval muscle; the *dpy-7* promoter, which is active in hypodermis; the *vha-6* promoter, which is active in intestine; and the ubiquitous *eft-3* promoter. We chose these promoters because they included many of the tissues that had *unc-14* driven expression of GFP (Figure 2B). As shown in Figure 3, only the ubiquitously-expressed *Peft-3::hif-1* rescued the lethality of *hif-1(ia04)* mutants exposed to H2S. Although we did not test all possible cell and tissue types, these data suggest that HIF-1 activity in a single tissue cannot support survival in H_2_S.

**Figure 3.**
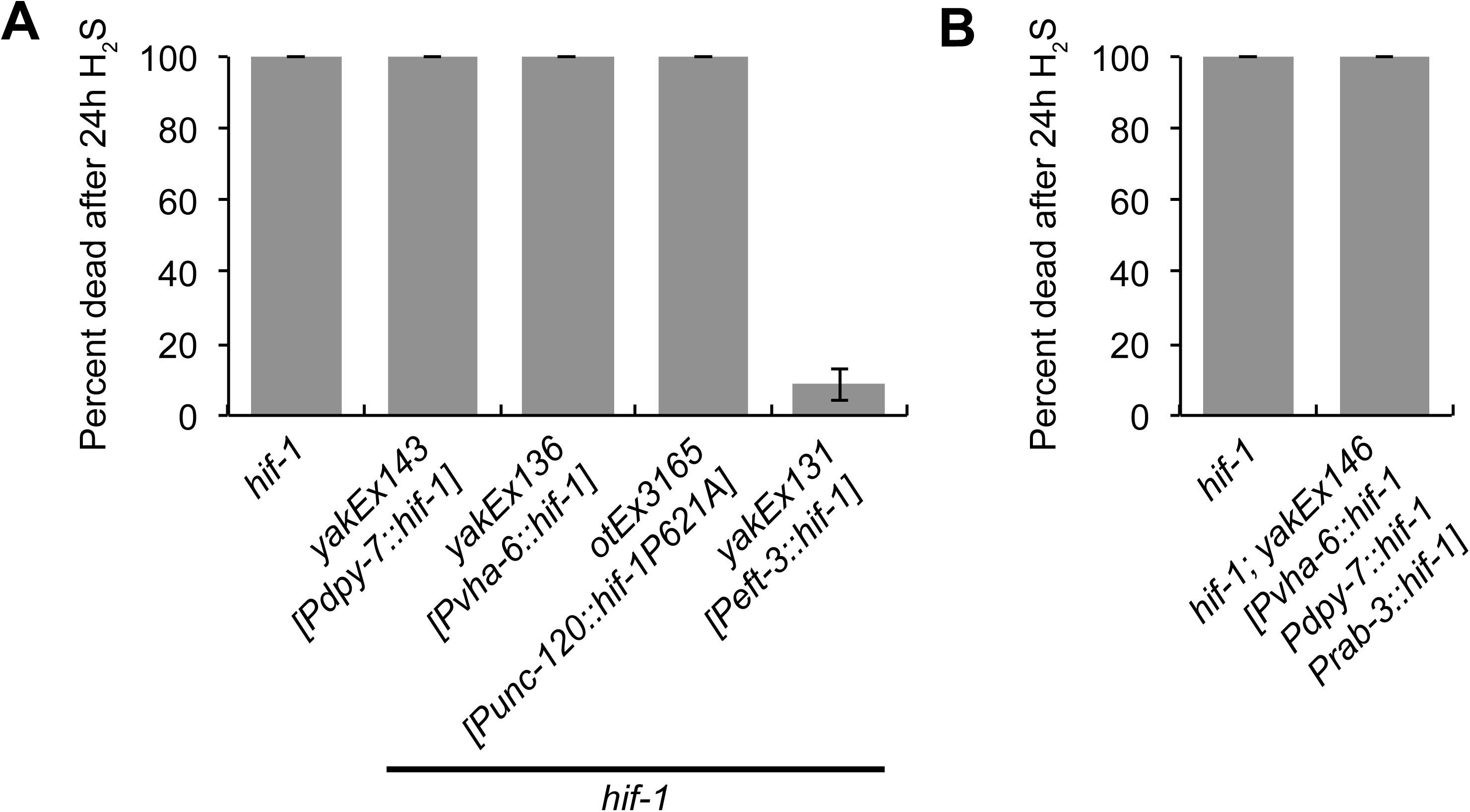
Survival in H_2_S requires broad expression of *hif-1*. Survival of animals exposed to H_2_S. All animals have the null *hif-1(ia04)* mutation. (A) Lethality of animals that express *hif-1* only in hypodermis (*Pdpy-7::hif-1; yakEx143*), intestine (*Pvha-6::hif-1; yakEx136*), or muscle (*Punc-120::hif-1(P621A); otEx3165*). As a control, *hif-1* was expressed from a ubiquitous promoter *(Peft-3::hif-1; yakEx131)*. Expression was from extrachromosomal arrays. Wild-type *hif-1* was used for all constructs except the *Punc-120::hif-1(P621A)*, which expresses the non-degradable variant. (B) Survival of *hif-1(ia04); yakEx146* animals exposed to H_2_S that express *hif-1* simultaneously in intestine (*Pvha-6::hif-1*), hypodermis (*Pdpy-7::hif-1*), and neurons (*Prab-3::hif-1*). Average of three independent experiments is shown, each with n = 20-35 animals. Error bars are standard error of the mean (SEM).

The fact that *Punc-14::hif-1* was sufficient for survival in H2S (Figure 1A) suggests that activity of HIF-1 may not be required in all cells. Since we did not observe rescue when *hif-1* was expressed in a single tissue, we made transgenic animals with expression of *hif-1* in more than one tissue to determine if we could find a minimal expression that was sufficient for survival in H_2_S. We found that even animals with *hif-1* expression in neurons, hypodermis, and intestine (*hif-1(ia04); yakEx146[Prab-3::hif-1, Pvha-6::hif-1, Pdpy-7::hif-1]*) did not survive exposure to H_2_S (Figure 3B). Together, our data suggests that that HIF-1 activity is required in many tissues to coordinate the essential response to H_2_S. This could indicate that HIF-1 acts cell autonomously to direct expression of many tissue-specific transcripts that are required to survive exposure to H_2_S.

Although it was reported that *otIs197* expresses *hif-1* selectively in neurons (Pocock and Hobert, 2008), our data show that the *unc-14* promoter is more broadly expressed. In fact, others have reported non-neuronal expression of transgenes expressed under the control of the *unc-14* promoter (Ogura *et al.*, 1997; Wolkow *et al.*, 2000; da Graca *et al.*, 2004). However, the non-neuronal expression we have demonstrated is much more penetrant than has been previously acknowledged. This is an important consideration when interpreting the results of experiments using transgenes driven by *unc-14*, including *hif-1* from *otIs197*. Our data show that non-neuronal expression from the *unc-14* promoter is significant and that rescue by *unc-14*-driven transgenes is not sufficient to infer neuronal function of HIF-1 and, presumably, other proteins.

## ACKNOWLEDGEMENTS

We would like to thank Dr. Michael Ailion (University of Washington) for sharing plasmids, discussing ideas, helping identify the different *C. elegans* tissues, and providing useful feedback on drafts of this manuscript. We are also grateful to Dr. Roger Pocock (Monash University) and Dr. Oliver Hobert (Columbia University) for sharing plasmids and strains. We thank Dr. Suzanne Hoppins (University of Washington) and Dr. Andrea Wills (University of Washington) for critical reading of the manuscript. Some strains were provided by the CGC, which is funded by NIH Office of Research Infrastructure Programs (P40 OD010440). This work was supported by NIH grant R01 ES024958 to DLM. DLM is a New Scholar in Aging of the Ellison Medical Foundation.

